# Driving potent neutralization of a SARS-CoV-2 Variant of Concern with a heterotypic boost

**DOI:** 10.1101/2021.04.03.438330

**Authors:** Daniel J. Sheward, Marco Mandolesi, Egon Urgard, Changil Kim, Leo Hanke, Laura Perez Vidakovics, Alec Pankow, Natalie L. Smith, Xaquin Castro Dopico, Gerald McInerney, Jonathan M. Coquet, Gunilla B. Karlsson Hedestam, Ben Murrell

## Abstract

The emergence of SARS-CoV-2 Variants of Concern (VOCs) with mutations in key neutralizing antibody epitopes threatens to undermine vaccines developed against the pandemic founder variant (Wu-Hu-1). Widespread vaccine rollout and continued transmission are creating a population that has antibody responses of varying potency to Wu-Hu-1. Against this background, it is critical to assess the outcomes of subsequent immunization with variant antigens. It is not yet known whether heterotypic vaccine boosts would be compromised by original antigenic sin, where pre-existing responses to a prior variant dampen responses to a new one, or whether the primed memory B cell repertoire would bridge the gap between Wu-Hu-1 and VOCs. Here, we show that a single adjuvanted dose of receptor binding domain (RBD) protein from VOC 501Y.V2 (B.1.351) drives an extremely potent neutralizing antibody response capable of cross-neutralizing both Wu-Hu-1 and 501Y.V2 in rhesus macaques previously immunized with Wu-Hu-1 spike protein. Passive immunization with plasma sampled following this boost protected K18-hACE2 mice from lethal challenge with a 501Y.V2 clinical isolate, whereas only partial protection was afforded by plasma sampled after two Wu-Hu-1 spike immunizations.

## Introduction

At least 20 candidate SARS-CoV-2 vaccines have already entered phase 3 clinical trials. A number of these demonstrated high efficacy ^1–5^, significantly reducing morbidity and mortality, and are being rolled-out globally. This first generation of vaccines all encode or deliver a spike glycoprotein derived from the pandemic founder strain, Wu-Hu-1^6^.

Driven by multiple evolutionary forces ^8^, SARS-CoV-2 is evading immune responses and undermining our prevention and mitigation strategies. Globally, a number of VOCs are rising in frequency (see Fig 1), each harbouring spike mutations that confer resistance to prior immunity. Of particular concern is the surge of variant 501Y.V2 ^9^, with multiple mutations in dominant neutralizing antibody epitopes making it several fold more resistant to antibodies elicited by current vaccines ^10–12^. This underpins the substantially reduced vaccine efficacies in South Africa, where this variant is circulating at high frequency ^13,14^. Updated vaccines are likely required to protect against current and future mutated variants. Importantly, by the time these are rolled out, a significant proportion of the global population are likely to be seropositive as a result of either infection, or immunization with Wu-Hu-1 based vaccines. A relevant question now is whether a single dose will be sufficient to induce robust neutralising antibody responses to VOCs in seropositive individuals, and whether these boosts are sufficient to confer protection. Importantly, the first exposure to a pathogen can shape future responses to mutated variants. This immunological imprinting or original antigenic sin ^15^ is well-described for influenza A virus, where protection is highest against the first strain encountered and diminished against those encountered later in life ^16,17^. It is crucial for the design of updated vaccines and regimens to determine if existing immunity dampens antibody responses to new VOCs, or if a heterotypic boost can efficiently recruit cross-protective memory responses.

**Fig. 1.**
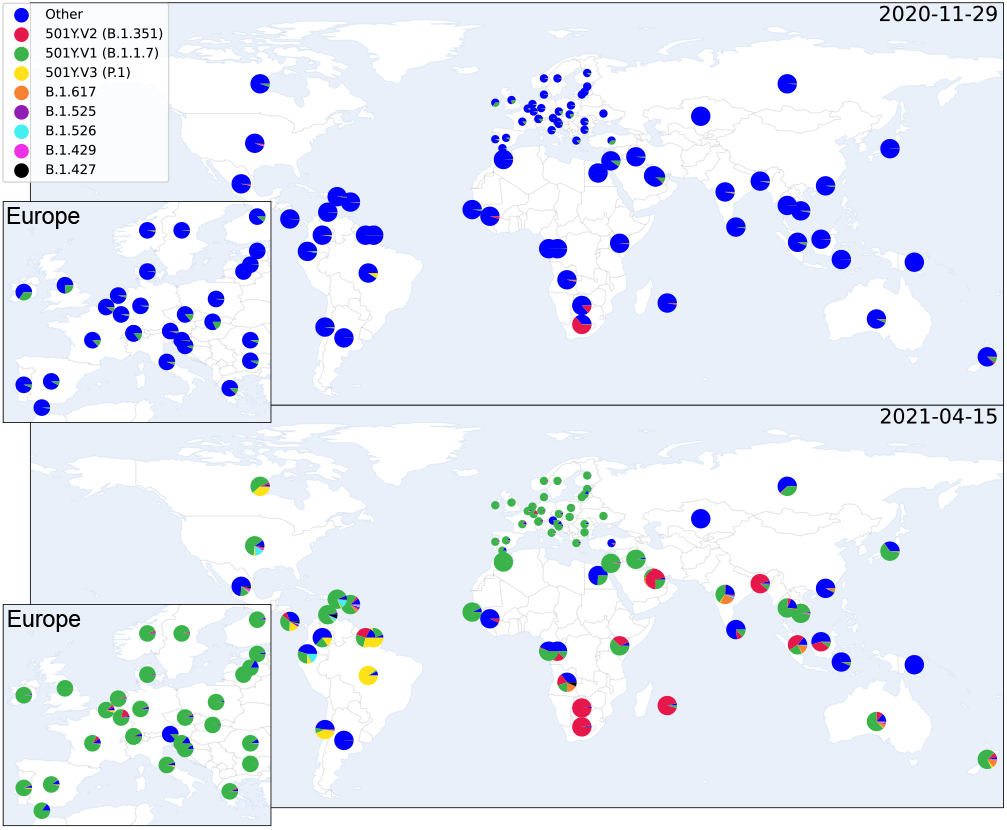
SARS-CoV-2 variants are rapidly coming to dominate the global genomic landscape. The global distribution and estimated country-level proportions of deposited SARS-CoV-2 genomes for eight variants, shown for 29th Nov 2020 (top), and five and a half months later for 15th April 2021 (bottom). Proportions over time are estimated from GISAID ^7^ genome metadata, using a temporally non-linear multinomial regression model (see methods).

## Results

To address this, we immunized three rhesus macaques with two doses of soluble prefusion-stabilized Wu-Hu-1 spike protein (2 µg), adjuvanted with saponin-based Matrix-M™ (Novavax AB, Uppsala, Sweden), with a one-month interval between doses, mimicking an immunization schedule for approved SARS-CoV-2 vaccines. After a single dose, neutralizing antibodies were detectable against Wu-Hu-1 but not against 501Y.V2 (Fig. 2). Neutralizing antibody responses against Wu-Hu-1 were substantially boosted by the second immunization (“post S”, GMT = 3980 at peak), and then waned over the following months (Fig. 2), as also reported in immunized humans ^19^. Notably, VOC 501Y.V2 was on average 9-fold (range: 5.6 - 12.2 fold) less potently neutralized (GMT = 451 at peak), consistent with the responses observed in humans following vaccination ^10–12^.

**Fig. 2.**
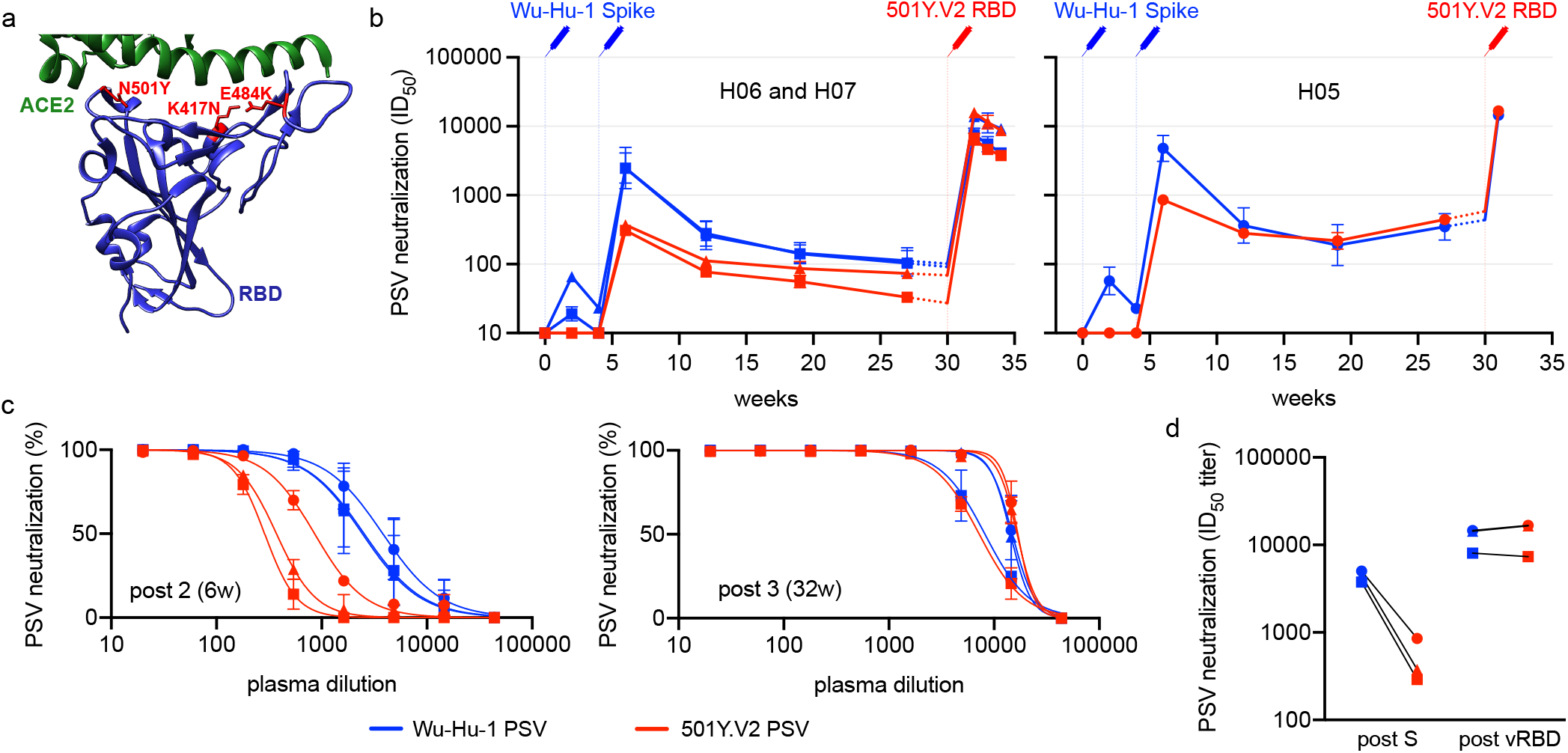
Heterotypic RBD boost drives a potent cross-neutralizing antibody response. (a) Depiction of the RBD immunogen (PDB:6MOJ ^18^) used as a heterotypic boost in this study, that incorporates the three RBD mutations (located in red) defining lineage 20H/501Y.V2. The cellular receptor, ACE2, is shown in green. (b-d) Neutralizing antibody responses over time to Wu-Hu-1 (blue) and 501Y.V2 (red) pseudotyped viruses (PSV) are shown for three immunized macaques: (b) H06 and H07 (left) and H05 (right), plotted separately as they exhibit different trajectories prior to the heterotypic boost. Syringes indicate the timing of immunizations (blue: Wu-Hu-1 spike at 0 and 4-weeks, red: 501Y.V2 RBD at 30-weeks). Titers from 27-30 weeks (shown with dashed lines) have been extrapolated for clarity. Error bars depict the geometric SD. (c) While neutralization of 501Y.V2 was significantly reduced at 6 weeks, corresponding to peak responses 2 weeks following the second spike dose (left), neutralization was restored following subsequent heterotypic RBD boost (right), such that 501Y.V2 (red) and Wu-Hu-1 (blue) were potently neutralized at similar titers (d) in all three animals.

Six months after their first immunization, macaques were boosted with either 2 µg (H05), 10 µg (H06), or 50 µg (H07) of soluble 501Y.V2 RBD in 50 µg Matrix-M™ adjuvant. One macaque (H05) was terminated 5 days after immunization, due to an unrelated illness that had begun prior to the third immunization, and was sampled for detailed follow-up studies of antibody specificities. The two other macaques (H06 and H07) were followed for 4 weeks. In all three animals, 501Y.V2 RBD efficiently boosted responses that potently cross-neutralized both Wu-Hu-1 and 501Y.V2, with similar titers (Fig. 2a-c; Wu-Hu-1 GMT = 11795, 501Y.V2 GMT = 12595). In contrast, for macaques previously immunized with three doses of Wu-Hu-1 spike ^20^, the reduced neutralization of 501Y.V2 compared to Wu-Hu-1 remained after the third homotypic spike immunization (Supp. Fig. 1).

To determine whether restoration of neutralizing antibody titers to 501Y.V2 afforded a biologically relevant improvement in protective immunity, mice transgenic for human ACE2 (K18-hACE2^21^) were passively immunized intraperitoneally (i.p.) with plasma samples taken either 2 weeks following the second spike immunization (N=8) (“post S”), or 1-2 weeks following the RBD booster immunization (N=8) (“post vRBD”). Passive immunization conferred titers approximately 10-fold lower than donor plasma (Supp. Fig 2), and macaque polyclonal antibodies were not rapidly cleared following xenotransfusion with an unchallenged mouse still maintaining titers >1400 after 5 days (data not shown). Mice were then challenged intranasally with 2.4×106 RNA copies of either 501Y.V2 or ‘wild-type’ (encoding a spike matching Wu-Hu-1) virus (corresponding to 100 PFU of 501Y.V2 or 86 PFU of wild-type), and weight — a reliable proxy for disease severity ^22^ — was monitored daily.

Across all groups, protection was strongly correlated with the neutralizing antibody titers to the challenge virus on the day of challenge (Spearman’s ρ = 0.822, p<1×10-8, Fig. 3a). All control mice that did not receive plasma (PBS only) succumbed to disease when challenged with either variant, showing precipitous weight loss starting around three days post-challenge (Fig. 3b-d). Passive transfer of post S serum conferred protection from WT virus (Fig. 3c) but not from 501Y.V2 (Fig. 3d), clearly demonstrating that evasion of the antibody response by this VOC was sufficient to cause disease. Notably, passive transfer of post vRBD plasma protected against both WT and 501Y.V2 (Fig. 3c,d).

**Fig. 3.**
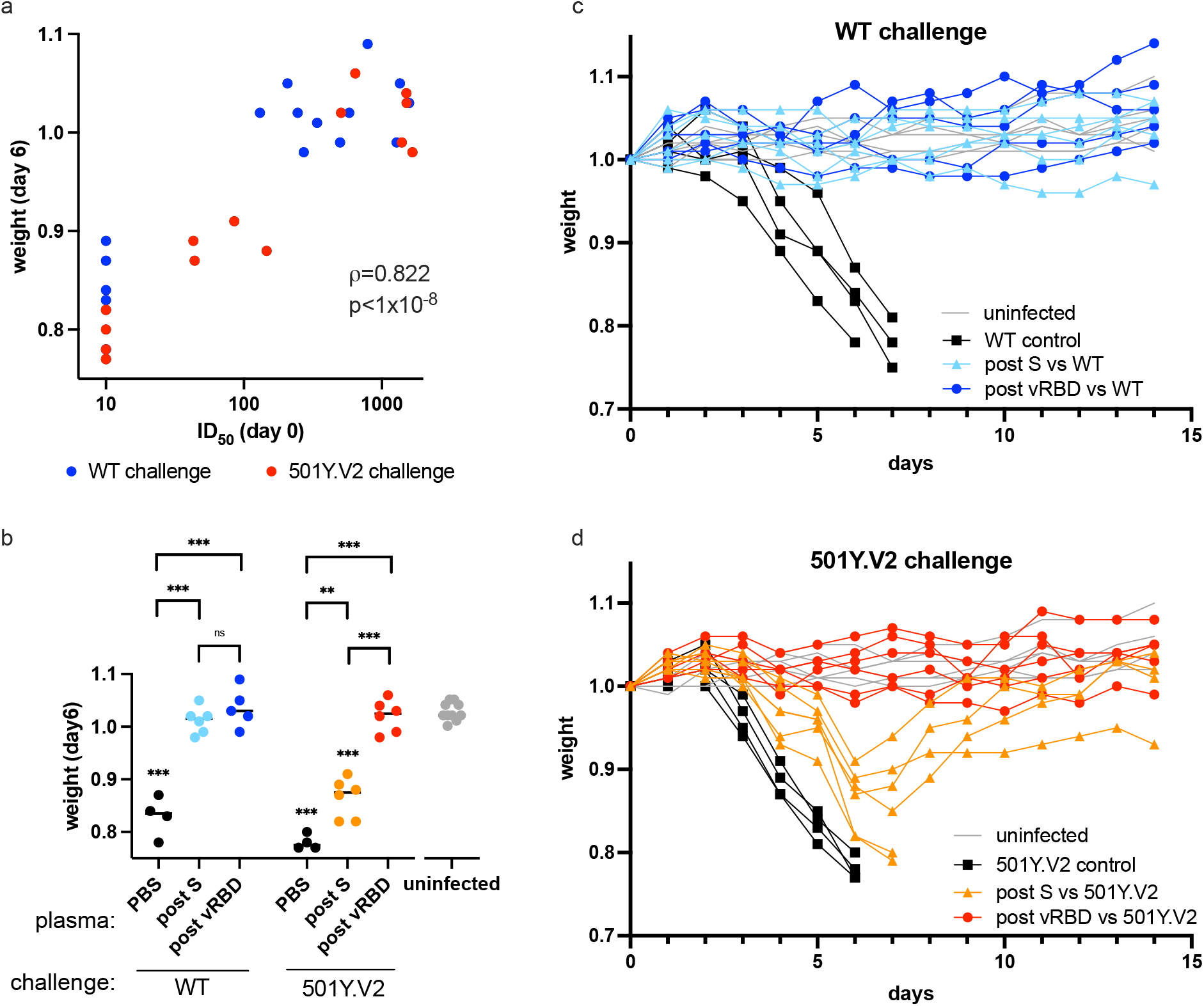
Heterotypic RBD boost restores protection against 501Y.V2 in passively immunized k18-hACE2 mice. (a) Pseudovirus neutralizing antibody titers against the challenge spike (ID_50_) in passively immunized mice on the day of challenge are associated with infection, and disease severity summarized as weight loss 6 days following challenge. Titers below the limit of detection of the assay (20) are plotted as 10. (b) Weight loss at day 6 for each group. Unchallenged littermates housed in the same cages (grey); PBS, mock immunized mice (black). Post S, passive immunization with plasma following the second spike immunization (6 week plasma); post vRBD, passive immunization with plasma from macaques boosted with variant (501Y.V2) RBD (31 or 32 week plasma). Statistical comparisons are summarized as: **, p*≤*0.01; * * *, p*≤*0.001 ; ns, not significant. Groups displaying significant weight loss compared to uninfected mice are annotated above the points for that group. (c-d) Weight loss following challenge with either (c) ‘wild-type’ (“WT”) or (d) 501Y.V2 virus for K18-hACE2 mice passively immunized with NHP plasma sampled post S or post vRBD. Control mice mock immunized with PBS and subsequently challenged (“PBS”) are shown in black, while uninfected littermates housed in the same cages (“uninfected”) are shown in grey.

## Discussion

For many licensed vaccines, reduced efficacy has been observed against the 501Y.V2 variant ^13,14,23^. Moreover, the decay of vaccine-elicited antibody titers ^19,24^ suggests that over time, protection will wane further. Consistent with reduced vaccine efficacy against 501Y.V2, we show that breakthrough infection and disease occurs in K18-hACE2 mice passively immunized with plasma from rhesus macaques immunized with two doses of adjuvanted wild-type spike protein.

Cross-species passive immunization and challenge experiments have been used to show antibody-mediated protection conferred by influenza vaccination (human to mouse ^25^), and more recently, to characterize protection from wild-type SARS-CoV-2 (macaque to hamster ^26^). Our data further demonstrate the utility of this approach for investigating protection against SARS-CoV-2 variants.

The ability of vaccines to broaden existing responses to new variants is still largely unclear. Despite weak immunogenicity of soluble, monomeric RBD as a priming antigen ^20^, heterotypic RBD administered as a boost elicited a potent recall response in non-human primates. This was robust to the boosting dose, and effective as low as 2 µg, possibly aided by a dose-sparing effect of Matrix-M ^27^. While reduced neutralization of 501Y.V2 was evident following 2 doses of Wu-Hu-1 spike, both Wu-Hu-1 and 501Y.V2 were potently neutralized following heterotypic (501Y.V2) RBD boost. In animal challenge models, neutralizing antibodies following passive immunization represented a robust correlate of protection such that the restoration of neutralizing antibody titers to 501Y.V2 also translated into protective immunity.

The potent, cross-neutralizing antibody response that arises following a heterotypic boost indicates that original antigenic sin does not represent a significant barrier to the acquisition of protective immunity against current SARS-CoV-2 VOCs. In the immunized animal sampled only 5 days post vRBD boost, neutralizing titers (against both Wu-Hu-1 and 501Y.V2) were already elevated suggesting these titers are the product of a rapidly activated population of antibody secreting cells. Further, this time course indicates that successive rounds of affinity maturation likely were not required for neutralization of 501Y.V2, but rather that vRBD-specific antibody responses could be boosted from the pool of existing memory B cells primed by Wu-Hu-1. These responses are largely consistent with recently reported results from 501Y.V2 spike mRNA (mRNA1273.351, Moderna) booster vaccinations ^28,29^. The observation that immunization with RBD (and not whole spike) was capable of inducing robust neutralising antibody responses is particularly promising as RBD is a small, stable protein that can be rapidly synthesized and efficiently expressed.

Taken together, these data indicate that potent, cross-neutralizing and cross-protective antibody responses can be recruited with heterotypic SARS-CoV-2 immunogens following a primary exposure, and identify soluble RBD booster immunizations as an attractive strategy to broaden vaccine protection from new SARS-CoV-2 variants.

## ACKNOWLEDGEMENTS

We thank Dr. Bengt Eriksson and all personnel at Astrid Fagraeus laboratory for expert assistance with rhesus macaques. We thank Monika Àdori for assistance processing samples. We also thank Novavax, AB, Uppsala, Sweden, for generously making the Matrix-M™ adjuvant available. We gratefully thank James Voss, Deli Huang, and Jesse Bloom for reagents. We gratefully acknowledge Penny Moore and the NICD (South Africa) for providing the 501Y.V2 spike plasmid (used here for PSV neutralization assays) which was generated using funding from the South African Medical Research Council. We thank Jonas Klingstrom for sharing the Swedish SARS-CoV-2 isolate and Alex Sigal from the Africa Health Research Institute for providing the 501Y.V2 isolate. For SARS-CoV-2 variant data from GISAID, we gratefully acknowledge all data contributors i.e. the Authors and their Originating Laboratories responsible for obtaining the specimens, and their Submitting Laboratories that generated the genetic sequence and metadata and shared via the GISAID Initiative the data on which the variant frequency estimates in this research are based. This project has received funding from the European Union’s Horizon 2020 research and innovation programme under grant agreement No. 101003653 (CoroNAb), to GM, GKH, and BM, from the Swedish Research Council to GM, JMC, GKH, and BM, and from Karolinska Institutet Development Office and Karolinska Institutet President’s Fund to GM, GKH, and BM. EU is supported by a Wenner Gren Fellowship.

## AUTHOR CONTRIBUTIONS

Conceptualization: DJS, MM, EU, JMC, GKH, BM; Formal Analysis: DJS, BM; Funding acquisition: GMM, JMC, GKH, BM; Investigation: DJS, MM, EU, CK, AP, NLS, XDC; Methodology: DJS, MM, EU, AP, NLS, JMC, GKH, BM; Resources: CK, LH, LPV; Software: BM; Supervision: LH, GMM, JMC, GKH, BM; Visualization: DJS, AP, BM; Writing – original draft: DJS, BM; Writing – review editing: all authors;

## METHODS

### Ethics statement

The animal work was conducted with the approval of Stockholms Jordbruksverket (10513-2020, 18427-2019 and 10895-2020). All animal procedures were performed according to approved guidelines.

### Animal models

Rhesus macaques (Macaca mulatta) of Chinese origin, 5-6 years old, were housed at the Astrid Fagraeus Laboratory at Karolinska Institutet. Housing and care procedures complied with the provisions and general guidelines of the Swedish Board of Agriculture. The facility has been assigned an Animal Welfare Assurance number by the Office of Laboratory Animal Welfare (OLAW) at the National Institutes of Health (NIH). The macaques were housed in groups in enriched 14 m3 cages. They were habituated to the housing conditions for more than six weeks before the start of the experiment and subjected to positive reinforcement training in order to reduce the stress associated with experimental procedures. The macaques were weighed at each sampling. All animals were confirmed negative for simian immunodeficiency virus, simian T cell lymphotropic virus, simian retrovirus type D and simian herpes B virus. Mice transgenic for human ACE2 under control of the cytokeratin 18 (K18) promoter ^21^ were obtained from the Jackson Laboratory. Mice were maintained as a hemizygous line, with hACE2 transgene presence confirmed using Sanger sequencing as per the Jackson Laboratory protocol. All mice were 12-21 weeks old at the start of the study, and experiments were conducted in BSL3 facilities at the Comparative Medicine department (KM-F) at Karolinska Institutet.

### Cell lines

HEK293T and HEK293T-hACE2 (Human, female) cells were cultured in a humidified 37°C incubator (5% CO2) in Dulbecco’s Modified Eagle Medium (Gibco) supplemented with 10% Fetal Bovine Serum and 1% Penicillin/Streptomycin, and were passaged when nearing confluency using 1X Trypsin-EDTA.

Calu-3 cells (Human, male), a lung-derived adenocarcinoma cell line, were obtained from Jonas Klingstrom (Karolinska Institutet). Calu-3 cells were maintained in Dulbecco’s Modified Eagle Medium supplemented with Ham’s F-12 (Thermo Fisher Scientific), 2% Fetal Bovine Serum and 1% Penicillin/Streptomycin in a humidified 37°C incubator (5% CO2) and were passaged using 0.5X Trypsin-EDTA.

Vero E6 cells (ATCC-CRL-1586, African Green Monkey) were maintained in DMEM (Gibco) supplemented with 2% fetal calf serum and 1% penicillin-streptomycin in a humidified incubator with 5% CO2 at 37°C.

## Method Details

### Immunizations

501Y.V2 RBD (encoding amino acid mutations K417N, E484K, and N501Y, and a C-terminal His-tag) was synthesized (Integrated DNA Technologies), and cloned into a mammalian expression vector (pcDNA3.1), using a Gibson Assembly Mastermix (New England Biolabs). Spike ectodomain (prefusion stabilized with 6 prolines ^30^) and RBD were produced by the transient transfection of Freestyle 293-F cells using FreeStyle MAX reagent (Thermo Fisher) or polyethylenimine (PEI), respectively. The HIS-tagged Spike ectodomain and RBD were purified from filtered supernatant using nickel IMAC resin (HisPur Ni-NTA, Thermo Fisher Scientific) followed by size-exclusion chromatography on a Superdex 200 (Cytiva) in PBS. On the day of immunization, indicated doses were mixed with 50 µg Matrix-M™ adjuvant (Novavax AB, Uppsala, Sweden) in a final inoculation volume of 800 µl.

Macaques were immunized intramuscularly (i.m.) with half of each dose administered in each quadricep. All immunizations and blood samplings were performed under sedation with 10-15 mg/kg ketamine (Ketaminol, Intervet, Sweden) administered i.m.. Blood plasma was isolated by centrifugation, and heat inactivated at 56°C for 60 minutes.

### Pseudotyped virus neutralization assays

All plasma and serum samples were heat inactivated at 56°C for 60 minutes. Pseudotyped lentiviruses displaying either the SARS-CoV-2 pandemic founder variant (Wu-Hu-1) or 501Y.V2 variant spike ^31^ and packaging a firefly luciferase reporter gene were generated by the co-transfection of HEK293T cells using Lipofectamine 3000 (Invitrogen) per the manufacturer’s protocols. Media was changed 12-16 hours after transfection, and pseudotyped viruses were harvested at 48- and 72-hours post-transfection, clarified by centrifugation, and stored at -80°C until use. Pseudotyped viruses sufficient to generate 50,000 relative light units (RLUs) were incubated with serial dilutions of plasma for 60 min at 37°C in a 96-well plate, and then 15,000 HEK293T-hACE2 cells were added to each well. Plates were incubated at 37°C for 48 hours, and luminescence was then measured using Bright-Glo (Promega) per the manufacturer’s protocol, on a GM-2000 luminometer (Promega).

### Virus production and quantification

Viral isolates expanded from clinical samples of ‘wild-type’ and 501Y.V2 ^32^ were kind gifts from Jonas Klingstrom (Karolinska Institutet) and Alex Sigal (The African Health Research Institute) respectively. Isolates were propagated in Calu-3 cells for 72 hours and harvested from the supernatant to generate replication competent SARS-CoV-2 stocks for animal challenge. All virus challenge stocks harboured no cell-culture adaptation mutations detectable by Sanger sequencing of the full-length spike gene (see Supp. Fig. 3). Viral titers were quantified by a plaque forming assay in Vero E6 cells, as previously described ^33^.

Viral RNA was quantified from clarified viral supernatant in TRIzol (ThermoFisher Scientific) and extracted using the manufacturer’s protocol but with the following modifications: Total RNA was precipitated with isopropanol in the presence of Glycoblue (Thermo Fisher Scientific) coprecipitant for 45 min at -20C, and RNA pellets were resuspended in warm RNase-free water.

RT-qPCR reactions were performed using the Superscript III one step RT-qPCR system with Platinum Taq Polymerase (Invitrogen) with 400 nM of each primer and 200 nM of probe. Primers and probes for the CoV-E gene target were as previously described ^34^. Thermal cycling consisted of RT at 55C for 10 min, denaturation at 95C for 3 min, and 45 cycles of 95C for 15 seconds and 58C for 30 seconds. Reactions were carried out using a CFX96 Connect Real-Time PCR Detection System (Bio-Rad) following manufacturer instructions. To generate standard curves, a synthetic DNA template gBlock (Integrated DNA Technologies) was transcribed using the mMessage mMachine T7 Transcription Kit (Invitrogen) and serially diluted. For each viral stock, RT-qPCR was run on a 10-fold dilution series ranging from 1:10 to 1:10,000 to ensure linearity of the assay and avoid saturation at high copy numbers.

### K18-hACE2 mouse challenge

One day prior to challenge, K18-hACE2 mice were passively immunized with 200µl of macaque plasma, administered intraperitoneally (i.p.) under isoflurane sedation. Each plasma sample was administered to four mice (two to be challenged with wild-type virus, and two to be challenged with 501Y.V2). The following day, mice were bled from the tail vein (to obtain serum for the characterization of neutralizing antibody titers), moved to a BSL3 facility and challenged intranasally with a standardized dose (2.4×106 RNA copies) of either ‘wild-type’ or 501Y.V2 virus stock in a total challenge volume of 40µl in PBS. All challenges were performed under light isoflurane sedation. Weight and general body condition was monitored daily until weight loss was evident, after which mice were monitored twice daily. Throughout the experiment, weight loss, changes in general health, breathing, body movement and posture, piloerection and eye health were monitored. Mice were euthanized when they experienced weight loss of at least 20% of their starting body weight, or when movement was greatly impaired and/or they experienced difficulty breathing that was considered to reach a severity level of 0.5 on Karolinska Institutet’s veterinary plan for monitoring animal health.

### Quantification and Statistical Analysis

#### Non-linear Multinomial Regression for VOC frequency estimation

SARS-CoV-2 lineage metadata was obtained from GISAID (gisaid.org - 2021-05-07 meta-data release) comprising 1,446,045 genomes. For each of the lineages in Figure 1, we aggregated daily counts of genomes at the country level, requiring at least 30 samples in the 30 days before 15th Feb, 2020, which was chosen because sequence data diminished rapidly beyond this point. Using a Generalized Linear Model, we model the daily variant counts with a multinomial distribution (and a log-link function), with underlying frequencies parameterized by a linear combination of 400 randomly drawn Fourier basis features (aka. a “Random Kitchen Sink” ^35^) to allow frequencies to vary non-linearly as a function of time. We estimate the model parameters with an L2 norm on the random feature coefficients, using the GLMNet.jl Julia package, plotting the map with Cartopy (https://github.com/SciTools/cartopy). Code available at https://github.com/MurrellGroup/VOCfreq.

#### ID50 titers

Neutralizing antibody ID50 titers were calculated in Prism 9 (GraphPad Software) by fitting a four-parameter logistic curve bounded between 0 and 100, and interpolating the reciprocal plasma/serum dilution where RLUs were reduced by 50% relative to control wells in the absence of plasma/serum.

### Statistical Analysis

Differences in day 6 weights between groups were assessed using Mann-Whitney tests in Prism 9 (Graphpad Software). Depicted symbols summarize the following: ns = p > 0.05 (not significant), * p *≤*0.05, ** p *≤* 0.01, *** p *≤* 0.001. The association between neutralizing antibody titers and weight loss was assessed with a Spearman’s rank correlation in Prism 9 (Graphpad Software).

---

**Fig. SI1.**
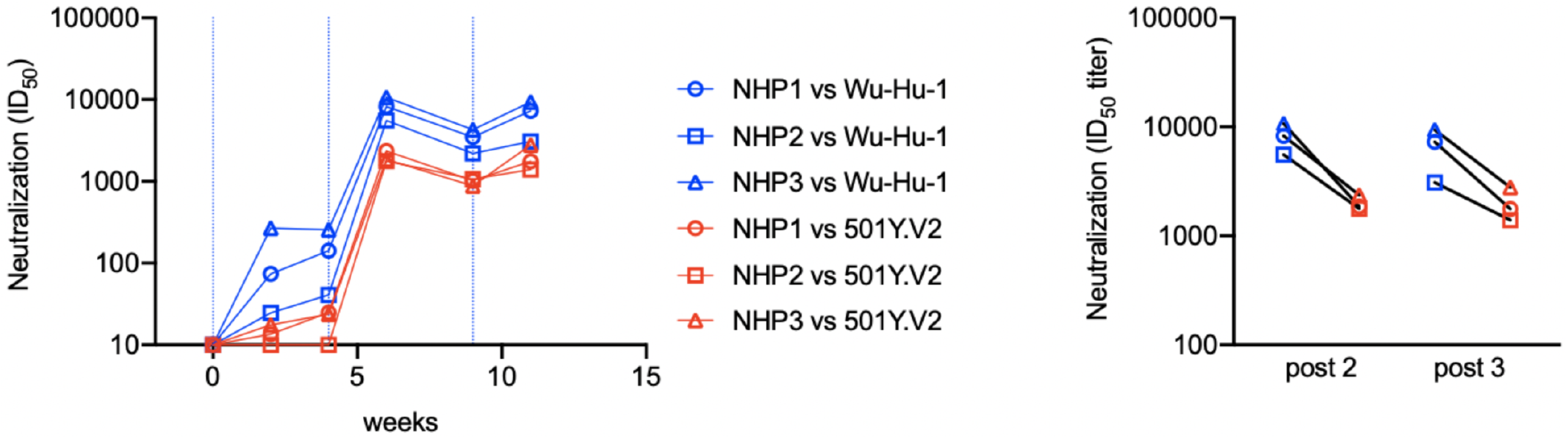
(left) Longitudinal neutralizing antibody responses against Wu-Hu-1 (blue) and 501Y.V2 (red) for plasma samples from Mandolesi et al. ^20^, where three rhesus macaques (NHP1-NHP3) were immunized with three doses of Wu-Hu-1 spike (100 µg) in Matrix-M™ adjuvant. Vertical blue lines indicate the timing of immunizations (at 0, 4, and 9 weeks). (right) Comparison of the titers at 6 weeks (post 2) and 11 weeks (post 3) illustrating that reduced titers to 501Y.V2 (red) compared to Wu-Hu-1 (blue) were maintained after a third homotypic spike boost.

**Fig. SI2.**
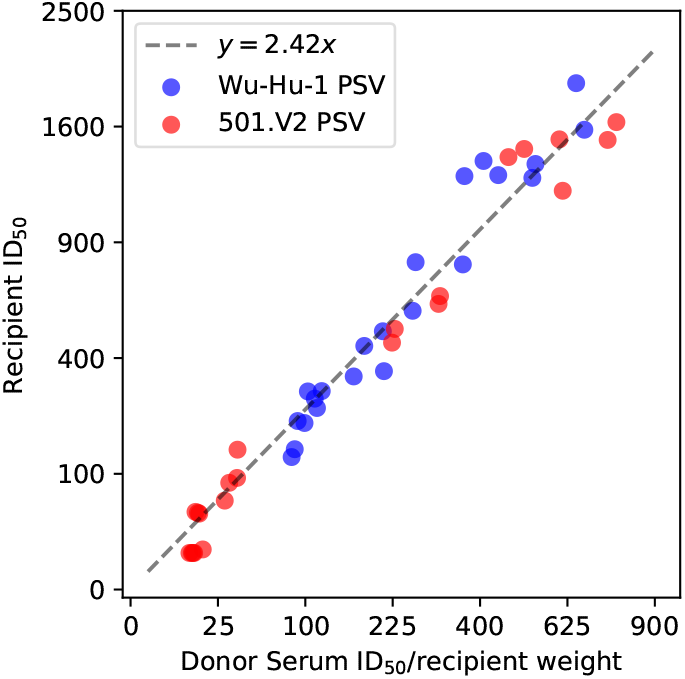
Relationship between donor and recipient titers following passive immunization. Mice were passively immunized with 200 ul of plasma, resulting in titers approximately 10-fold lower than in the donor. Correcting for weight, Spearman’s *r* = 0.98 (after a variance-stabilizing square-root transform).

**Fig. SI3.**
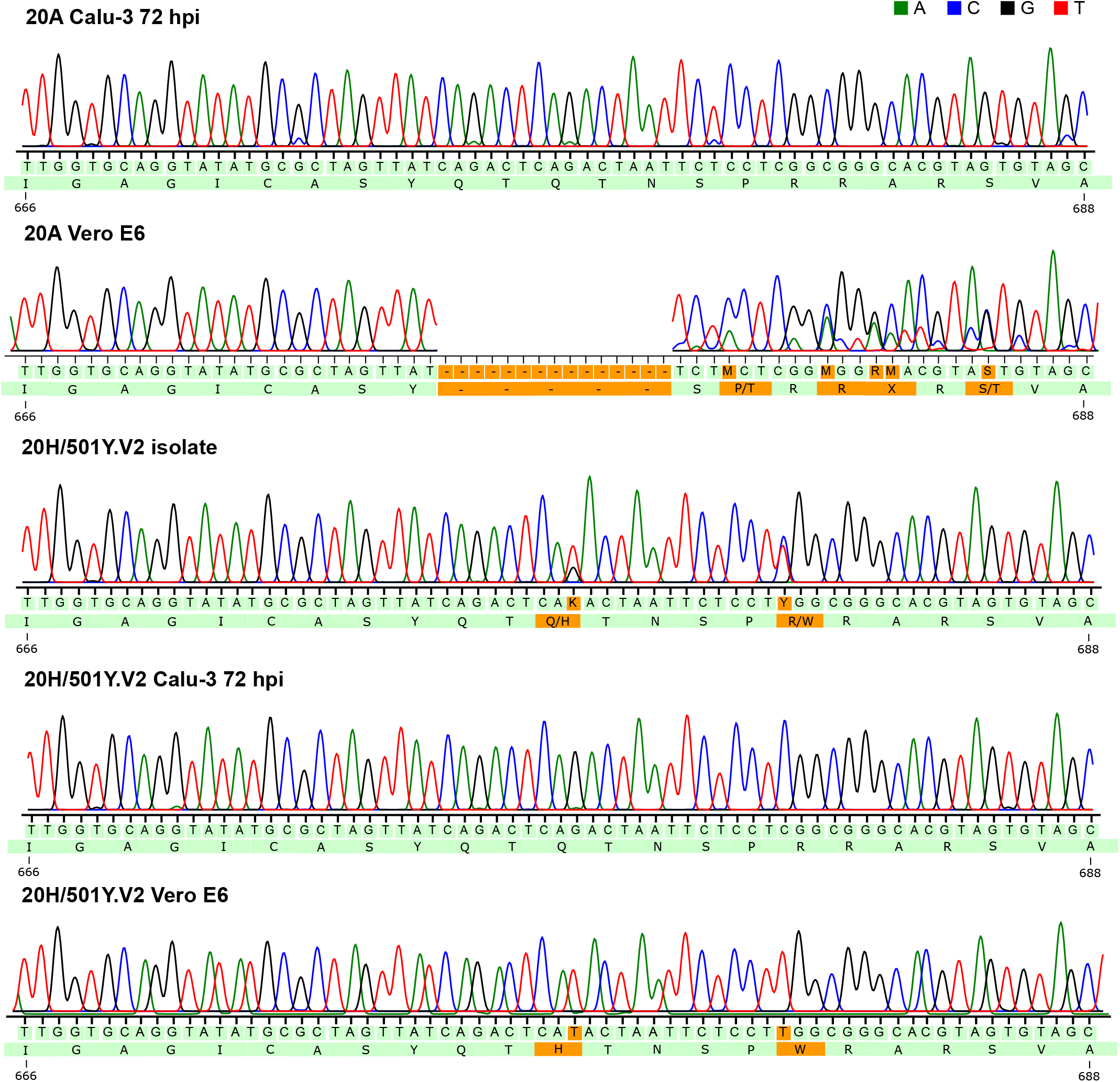
Expansion of SARS-CoV-2 isolates in Vero E6 but not Calu-3 cells rapidly selected for mutations and deletions proximal to the furin cleavage site. Challenge stocks used in this study were produced in Calu-3 cells, and confirmed by Sanger sequencing to harbour no high frequency cell culture adaptation mutations in spike. Electropherograms spanning the furin cleavage site from sanger sequencing of amplified viral RNA are shown for virus cultured in Vero E6 or Calu-3 cells demonstrating the rapid loss of the furin recognition sequence upon culture in Vero E6 cells but not Calu3 cells. Received stock of 501Y.V2 (“20H/501Y.V2 isolate”) had a mixture of intact/knocked-out furin site.

## Bibliography

1. Lindsey R Baden, Hana M El Sahly, Brandon Essink, Karen Kotloff, Sharon Frey, Rick Novak, David Diemert, Stephen A Spector, Nadine Rouphael, C Buddy Creech, John McGettigan, Shishir Khetan, Nathan Segall, Joel Solis, Adam Brosz, Carlos Fierro, Howard Schwartz, Kathleen Neuzil, Larry Corey, Peter Gilbert, Holly Janes, Dean Follmann, Mary Marovich, John Mascola, Laura Polakowski, Julie Ledgerwood, Barney S Graham, Hamilton Bennett, Rolando Pajon, Conor Knightly, Brett Leav, Weiping Deng, Honghong Zhou, Shu Han, Melanie Ivarsson, Jacqueline Miller, Tal Zaks, and COVE Study Group. Efficacy and safety of the mRNA-1273 SARS-CoV-2 vaccine. N. Engl. J. Med., 384(5):403–416, February 2021.

2. Fernando P Polack, Stephen J Thomas, Nicholas Kitchin, Judith Absalon, Alejandra Gurtman, Stephen Lockhart, John L Perez, Gonzalo Pérez Marc, Edson D Moreira, Cristiano Zerbini, Ruth Bailey, Kena A Swanson, Satrajit Roychoudhury, Kenneth Koury, Ping Li, Warren V Kalina, David Cooper, Robert W Frenck, Jr, Laura L Hammitt, Özlem Türeci, Haylene Nell, Axel Schaefer, Serhat Ünal, Dina B Tresnan, Susan Mather, Philip R Dormitzer, Uğ ur Ş ahin, Kathrin U Jansen, William C Gruber, and C4591001 Clinical Trial Group. Safety and efficacy of the BNT162b2 mRNA covid-19 vaccine. N. Engl. J. Med., December 2020.

3. Merryn Voysey, Sue Ann Costa Clemens, Shabir A Madhi, Lily Y Weckx, Pedro M Folegatti, Parvinder K Aley, Brian Angus, Vicky L Baillie, Shaun L Barnabas, Qasim E Bhorat, Sagida Bibi, Carmen Briner, Paola Cicconi, Andrea M Collins, Rachel Colin-Jones, Clare L Cutland, Thomas C Darton, Keertan Dheda, Christopher J A Duncan, Katherine R W Emary, Katie J Ewer, Lee Fairlie, Saul N Faust, Shuo Feng, Daniela M Ferreira, Adam Finn, Anna L Goodman, Catherine M Green, Christopher A Green, Paul T Heath, Catherine Hill, Helen Hill, Ian Hirsch, Susanne H C Hodgson, Alane Izu, Susan Jackson, Daniel Jenkin, Carina C D Joe, Simon Kerridge, Anthonet Koen, Gaurav Kwatra, Rajeka Lazarus, Alison M Lawrie, Alice Lelliott, Vincenzo Libri, Patrick J Lillie, Raburn Mallory, Ana V A Mendes, Eveline P Milan, Angela M Minassian, Alastair McGregor, Hazel Morrison, Yama F Mujadidi, Anusha Nana, Peter J O’Reilly, Sherman D Padayachee, Ana Pittella, Emma Plested, Katrina M Pollock, Maheshi N Ramasamy, Sarah Rhead, Alexandre V Schwarzbold, Nisha Singh, Andrew Smith, Rinn Song, Matthew D Snape, Eduardo Sprinz, Rebecca K Sutherland, Richard Tarrant, Emma C Thomson, M Estée Török, Mark Toshner, David P J Turner, Johan Vekemans, Tonya L Villafana, Marion E E Watson, Christopher J Williams, Alexander D Douglas, Adrian V S Hill, Teresa Lambe, Sarah C Gilbert, Andrew J Pollard, and Oxford COVID Vaccine Trial Group. Safety and efficacy of the ChAdOx1 nCoV-19 vaccine (AZD1222) against SARS-CoV-2: an interim analysis of four randomised controlled trials in brazil, south africa, and the UK. Lancet, December 2020.

4. Cheryl Keech, Gary Albert, Iksung Cho, Andreana Robertson, Patricia Reed, Susan Neal, Joyce S Plested, Mingzhu Zhu, Shane Cloney-Clark, Haixia Zhou, Gale Smith, Nita Patel, Matthew B Frieman, Robert E Haupt, James Logue, Marisa McGrath, Stuart Weston, Pedro A Piedra, Chinar Desai, Kathleen Callahan, Maggie Lewis, Patricia Price-Abbott, Neil Formica, Vivek Shinde, Louis Fries, Jason D Lickliter, Paul Griffin, Bethanie Wilkinson, and Gregory M Glenn. Phase 1–2 trial of a SARS-CoV-2 recombinant spike protein nanoparticle vaccine. N. Engl. J. Med., 383(24):2320–2332, December 2020.

5. Denis Y Logunov, Inna V Dolzhikova, Dmitry V Shcheblyakov, Amir I Tukhvatulin, Olga V Zubkova, Alina S Dzharullaeva, Anna V Kovyrshina, Nadezhda L Lubenets, Daria M Grousova, Alina S Erokhova, Andrei G Botikov, Fatima M Izhaeva, Olga Popova, Tatiana A Ozharovskaya, Ilias B Esmagambetov, Irina A Favorskaya, Denis I Zrelkin, Daria V Voronina, Dmitry N Shcherbinin, Alexander S Semikhin, Yana V Simakova, Elizaveta A Tokarskaya, Daria A Egorova, Maksim M Shmarov, Natalia A Nikitenko, Vladimir A Gushchin, Elena A Smolyarchuk, Sergey K Zyryanov, Sergei V Borisevich, Boris S Naroditsky, Alexander L Gintsburg, and Gam-COVID-Vac Vaccine Trial Group. Safety and efficacy of an rad26 and rad5 vector-based heterologous prime-boost COVID-19 vaccine: an interim analysis of a randomised controlled phase 3 trial in russia. Lancet, 397(10275):671–681, February 2021.

6. Fan Wu, Su Zhao, Bin Yu, Yan-Mei Chen, Wen Wang, Zhi-Gang Song, Yi Hu, Zhao-Wu Tao, Jun-Hua Tian, Yuan-Yuan Pei, Ming-Li Yuan, Yu-Ling Zhang, Fa-Hui Dai, Yi Liu, Qi-Min Wang, Jiao-Jiao Zheng, Lin Xu, Edward C Holmes, and Yong-Zhen Zhang. A new coronavirus associated with human respiratory disease in china. Nature, 579(7798):265–269, March 2020.

7. Yuelong Shu and John McCauley. GISAID: Global initiative on sharing all influenza data – from vision to reality, 2017.

8. Darren P Martin, Steven Weaver, Houryiah Tegally, Emmanuel James San, Stephen D Shank, Eduan Wilkinson, Jennifer Giandhari, Sureshnee Naidoo, Yeshnee Pillay, Lavanya Singh, Richard J Lessells, NGS-SA, COVID-19 Genomics UK (COG-UK), Ravindra K Gupta, Joel O Wertheim, Anton Nekturenko, Ben Murrell, Gordon W Harkins, Philippe Lemey, Oscar A MacLean, David L Robertson, Tulio de Oliveira, and Sergei L Kosakovsky Pond. The emergence and ongoing convergent evolution of the N501Y lineages coincides with a major global shift in the SARS-CoV-2 selective landscape. medRxiv, March 2021.

9. Houriiyah Tegally, Eduan Wilkinson, Marta Giovanetti, Arash Iranzadeh, Vagner Fonseca, Jennifer Giandhari, Deelan Doolabh, Sureshnee Pillay, Emmanuel James San, Nokukhanya Msomi, Koleka Mlisana, Anne von Gottberg, Sibongile Walaza, Mushal Allam, Arshad Ismail, Thabo Mohale, Allison J Glass, Susan Engelbrecht, Gert Van Zyl, Wolfgang Preiser, Francesco Petruccione, Alex Sigal, Diana Hardie, Gert Marais, Marvin Hsiao, Stephen Korsman, Mary-Ann Davies, Lynn Tyers, Innocent Mudau, Denis York, Caroline Maslo, Dominique Goedhals, Shareef Abrahams, Oluwakemi Laguda-Akingba, Arghavan Alisoltani-Dehkordi, Adam Godzik, Constantinos Kurt Wibmer, Bryan Trevor Sewell, José Lourenço, Luiz Carlos Junior Alcantara, Sergei L Kosakovsky Pond, Steven Weaver, Darren Martin, Richard J Lessells, Jinal N Bhiman, Carolyn Williamson, and Tulio de Oliveira. Emergence and rapid spread of a new severe acute respiratory syndrome-related coronavirus 2 (SARS-CoV-2) lineage with multiple spike mutations in south africa. December 2020.

10. Pengfei Wang, Manoj S Nair, Lihong Liu, Sho Iketani, Yang Luo, Yicheng Guo, Maple Wang, Jian Yu, Baoshan Zhang, Peter D Kwong, Barney S Graham, John R Mascola, Jennifer Y Chang, Michael T Yin, Magdalena Sobieszczyk, Christos A Kyratsous, Lawrence Shapiro, Zizhang Sheng, Yaoxing Huang, and David D Ho. Antibody resistance of SARS-CoV-2 variants b.1.351 and b.1.1.7. Nature, March 2021.

11. Daming Zhou, Wanwisa Dejnirattisai, Piyada Supasa, Chang Liu, Alexander J Mentzer, Helen M Ginn, Yuguang Zhao, Helen M E Duyvesteyn, Aekkachai Tuekprakhon, Rungtiwa Nutalai, Beibei Wang, Guido C Paesen, Cesar Lopez-Camacho, Jose Slon-Campos, Bassam Hallis, Naomi Coombes, Kevin Bewley, Sue Charlton, Thomas S Walter, Donal Skelly, Sheila F Lumley, Christina Dold, Robert Levin, Tao Dong, Andrew J Pollard, Julian C Knight, Derrick Crook, Teresa Lambe, Elizabeth Clutterbuck, Sagida Bibi, Amy Flaxman, Mustapha Bittaye, Sandra Belij-Rammerstorfer, Sarah Gilbert, William James, Miles W Carroll, Paul Klenerman, Eleanor Barnes, Susanna J Dunachie, Elizabeth E Fry, Juthathip Mongkolspaya, Jingshan Ren, David I Stuart, and Gavin R Screaton. Evidence of escape of SARS-CoV-2 variant b.1.351 from natural and vaccine induced sera. Cell, February 2021.

12. Kai Wu, Anne P Werner, Juan I Moliva, Matthew Koch, Angela Choi, Guillaume B E Stewart-Jones, Hamilton Bennett, Seyhan Boyoglu-Barnum, Wei Shi, Barney S Graham, Andrea Carfi, Kizzmekia S Corbett, Robert A Seder, and Darin K Edwards. mRNA-1273 vaccine induces neutralizing antibodies against spike mutants from global SARS-CoV-2 variants. bioRxiv, January 2021.

13. Shabir A Madhi, Vicky Baillie, Clare L Cutland, Merryn Voysey, Anthonet L Koen, Lee Fairlie, Sherman D Padayachee, Keertan Dheda, Shaun L Barnabas, Qasim E Bhorat, Carmen Briner, Gaurav Kwatra, Khatija Ahmed, Parvinder Aley, Sutika Bhikha, Jinal N Bhiman, As’ad E Bhorat, Jeanine du Plessis, Aliasgar Esmail, Marisa Groenewald, Elizea Horne, Shi-Hsia Hwa, Aylin Jose, Teresa Lambe, Matt Laubscher, Mookho Malahleha, Masebole Masenya, Mduduzi Masilela, Shakeel McKenzie, Kgaogelo Molapo, Andrew Moultrie, Suzette Oelofse, Faeezah Patel, Sureshnee Pillay, Sarah Rhead, Hylton Rodel, Lindie Rossouw, Carol Taoushanis, Houriiyah Tegally, Asha Thombrayil, Samuel van Eck, Constantinos K Wibmer, Nicholas M Durham, Elizabeth J Kelly, Tonya L Villafana, Sarah Gilbert, Andrew J Pollard, Tulio de Oliveira, Penny L Moore, Alex Sigal, Alane Izu, and NGS-SA Group Wits–VIDA COVID Group. Efficacy of the ChAdOx1 nCoV-19 covid-19 vaccine against the b.1.351 variant. N. Engl. J. Med., March 2021.

14. Vivek Shinde, Sutika Bhikha, Zaheer Hoosain, Moherndran Archary, Qasim Bhorat, Lee Fairlie, Umesh Lalloo, Mduduzi S L Masilela, Dhayendre Moodley, Sherika Hanley, Leon Fouche, Cheryl Louw, Michele Tameris, Nishanta Singh, Ameena Goga, Keertan Dheda, Coert Grobbelaar, Gertruida Kruger, Nazira Carrim-Ganey, Vicky Baillie, Tulio de Oliveira, Anthonet Lombard Koen, Johan J Lombaard, Rosie Mngqibisa, As’ad E Bhorat, Gabriella Benadé, Natasha Lalloo, Annah Pitsi, Pieter-Louis Vollgraaff, Angelique Luabeya, Aliasgar Esmail, Friedrich G Petrick, Aylin Oommen-Jose, Sharne Foulkes, Khatija Ahmed, Asha Thombrayil, Lou Fries, Shane Cloney-Clark, Mingzhu Zhu, Chijioke Bennett, Gary Albert, Emmanuel Faust, Joyce S Plested, Andreana Robertson, Susan Neal, Iksung Cho, Greg M Glenn, Filip Dubovsky, Shabir A Madhi, and 2019nCoV-501 Study Group. Efficacy of NVX-CoV2373 covid-19 vaccine against the b.1.351 variant. N. Engl. J. Med., May 2021.

15. Thomas Francis. On the doctrine of original antigenic sin. Proc. Am. Philos. Soc., 104(6): 572–578, 1960.

16. Justin Lessler, Steven Riley, Jonathan M Read, Shuying Wang, Huachen Zhu, Gavin J D Smith, Yi Guan, Chao Qiang Jiang, and Derek A T Cummings. Evidence for antigenic seniority in influenza a (H3N2) antibody responses in southern china. PLoS Pathog., 8(7):e1002802, July 2012.

17. Katelyn M Gostic, Monique Ambrose, Michael Worobey, and James O Lloyd-Smith. Potent protection against H5N1 and H7N9 influenza via childhood hemagglutinin imprinting. Science, 354(6313):722–726, November 2016.

18. Jun Lan, Jiwan Ge, Jinfang Yu, Sisi Shan, Huan Zhou, Shilong Fan, Qi Zhang, Xuanling Shi, Qisheng Wang, Linqi Zhang, and Xinquan Wang. Structure of the SARS-CoV-2 spike receptor-binding domain bound to the ACE2 receptor. Nature, 581(7807):215–220, May 2020.

19. Alicia T Widge, Nadine G Rouphael, Lisa A Jackson, Evan J Anderson, Paul C Roberts, Mamodikoe Makhene, James D Chappell, Mark R Denison, Laura J Stevens, Andrea J Pruijssers, Adrian B McDermott, Britta Flach, Bob C Lin, Nicole A Doria-Rose, Sijy O’Dell, Stephen D Schmidt, Kathleen M Neuzil, Hamilton Bennett, Brett Leav, Mat Makowski, Jim Albert, Kaitlyn Cross, Venkata-Viswanadh Edara, Katharine Floyd, Mehul S Suthar, Wendy Buchanan, Catherine J Luke, Julie E Ledgerwood, John R Mascola, Barney S Graham, John H Beigel, and mRNA-1273 Study Group. Durability of responses after SARS-CoV-2 mRNA-1273 vaccination. N. Engl. J. Med., December 2020.

20. Marco Mandolesi, Daniel J Sheward, Leo Hanke, Junjie Ma, Pradeepa Pushparaj, Laura Perez Vidakovics, Changil Kim, Monika Àdori, Klara Lenart, Karin Loré, Xaquin Castro Dopico, Jonathan M Coquet, Gerald M McInerney, Gunilla B Karlsson Hedestam, and Ben Murrell. SARS-CoV-2 protein subunit vaccination of mice and rhesus macaques elicits potent and durable neutralizing antibody responses. Cell Rep Med, 2(4):100252, April 2021.

21. Paul B McCray, Jr, Lecia Pewe, Christine Wohlford-Lenane, Melissa Hickey, Lori Manzel, Lei Shi, Jason Netland, Hong Peng Jia, Carmen Halabi, Curt D Sigmund, David K Meyerholz, Patricia Kirby, Dwight C Look, and Stanley Perlman. Lethal infection of K18-hACE2 mice infected with severe acute respiratory syndrome coronavirus. J. Virol., 81(2):813–821, January 2007.

22. Emma S Winkler, Adam L Bailey, Natasha M Kafai, Sharmila Nair, Broc T McCune, Jinsheng Yu, Julie M Fox, Rita E Chen, James T Earnest, Shamus P Keeler, Jon H Ritter, Liang-I Kang, Sarah Dort, Annette Robichaud, Richard Head, Michael J Holtzman, and Michael S Diamond. SARS-CoV-2 infection of human ACE2-transgenic mice causes severe lung inflammation and impaired function. Nat. Immunol., 21(11):1327–1335, November 2020.

23. Laith J Abu-Raddad, Hiam Chemaitelly, Adeel A Butt, and National Study Group for COVID-19 Vaccination. Effectiveness of the BNT162b2 covid-19 vaccine against the b.1.1.7 and b.1.351 variants. N. Engl. J. Med., May 2021.

24. Nicole Doria-Rose, Mehul S Suthar, Mat Makowski, Sarah O’Connell, Adrian B McDermott, Britta Flach, Julie E Ledgerwood, John R Mascola, Barney S Graham, Bob C Lin, Sijy O’Dell, Stephen D Schmidt, Alicia T Widge, Venkata-Viswanadh Edara, Evan J Anderson, Lilin Lai, Katharine Floyd, Nadine G Rouphael, Veronika Zarnitsyna, Paul C Roberts, Mamodikoe Makhene, Wendy Buchanan, Catherine J Luke, John H Beigel, Lisa A Jackson, Kathleen M Neuzil, Hamilton Bennett, Brett Leav, Jim Albert, Pratap Kunwar, and mRNA-1273 Study Group. Antibody persistence through 6 months after the second dose of mRNA-1273 vaccine for covid-19. N. Engl. J. Med., April 2021.

25. Raffael Nachbagauer, Jodi Feser, Abdollah Naficy, David I Bernstein, Jeffrey Guptill, Emmanuel B Walter, Franceso Berlanda-Scorza, Daniel Stadlbauer, Patrick C Wilson, Teresa Aydillo, Mohammad Amin Behzadi, Disha Bhavsar, Carly Bliss, Christina Capuano, Juan Manuel Carreño, Veronika Chromikova, Carine Claeys, Lynda Coughlan, Alec W Freyn, Christopher Gast, Andres Javier, Kaijun Jiang, Chiara Mariottini, Meagan McMahon, Monica McNeal, Alicia Solórzano, Shirin Strohmeier, Weina Sun, Marie Van der Wielen, Bruce L Innis, Adolfo García-Sastre, Peter Palese, and Florian Krammer. A chimeric hemagglutinin-based universal influenza virus vaccine approach induces broad and longlasting immunity in a randomized, placebo-controlled phase I trial. Nat. Med., 27(1):106–114, January 2021.

26. Kizzmekia S Corbett, Martha C Nason, Britta Flach, Matthew Gagne, Sarah O’ Connell, Timothy S Johnston, Shruti N Shah, Venkata Viswanadh Edara, Katharine Floyd, Lilin Lai, Charlene McDanal, Joseph R Francica, Barbara Flynn, Kai Wu, Angela Choi, Matthew Koch, Olubukola M Abiona, Anne P Werner, Gabriela S Alvarado, Shayne F Andrew, Mitzi M Donaldson, Jonathan Fintzi, Dillon R Flebbe, Evan Lamb, Amy T Noe, Saule T Nurmukhambetova, Samantha J Provost, Anthony Cook, Alan Dodson, Andrew Faudree, Jack Greenhouse, Swagata Kar, Laurent Pessaint, Maciel Porto, Katelyn Steingrebe, Daniel Valentin, Serge Zouantcha, Kevin W Bock, Mahnaz Minai, Bianca M Nagata, Juan I Moliva, Renee van de Wetering, Seyhan Boyoglu-Barnum, Kwanyee Leung, Wei Shi, Eun Sung Yang, Yi Zhang, John-Paul M Todd, Lingshu Wang, Hanne Andersen, Kathryn E Foulds, Darin K Edwards, John R Mascola, Ian N Moore, Mark G Lewis, Andrea Carfi, David Montefiori, Mehul S Suthar, Adrian McDermott, Nancy J Sullivan, Mario Roederer, Daniel C Douek, Barney S Graham, and Robert A Seder. Immune correlates of protection by mRNA-1273 immunization against SARS-CoV-2 infection in nonhuman primates. bioRxiv, April 2021.

27. Jing-Hui Tian, Nita Patel, Robert Haupt, Haixia Zhou, Stuart Weston, Holly Hammond, James Logue, Alyse D Portnoff, James Norton, Mimi Guebre-Xabier, Bin Zhou, Kelsey Jacobson, Sonia Maciejewski, Rafia Khatoon, Malgorzata Wisniewska, Will Moffitt, Stefanie Kluepfel-Stahl, Betty Ekechukwu, James Papin, Sarathi Boddapati, C Jason Wong, Pedro A Piedra, Matthew B Frieman, Michael J Massare, Louis Fries, Karin Lövgren Bengtsson, Linda Stertman, Larry Ellingsworth, Gregory Glenn, and Gale Smith. SARS-CoV-2 spike glycoprotein vaccine candidate NVX-CoV2373 immunogenicity in baboons and protection in mice. Nat. Commun., 12(1):372, January 2021.

28. Kai Wu, Angela Choi, Matthew Koch, Sayda Elbashir, Lingzhi Ma, Diana Lee, Angela Woods, Carole Henry, Charis Palandjian, Anna Hill, Julian Quinones, Naveen Nunna, Sarah O’Connell, Adrian B McDermott, Samantha Falcone, Elisabeth Narayanan, Tonya Colpitts, Hamilton Bennett, Kizzmekia S Corbett, Robert Seder, Barney S Graham, Guillaume B E Stewart-Jones, Andrea Carfi, and Darin K Edwards. Variant SARS-CoV-2 mRNA vaccines confer broad neutralization as primary or booster series in mice. April 2021.

29. Kai Wu, Angela Choi, Matthew Koch, Lingzhi Ma, Anna Hill, Naveen Nunna, Wenmei Huang, Judy Oestreicher, Tonya Colpitts, Hamilton Bennett, Holly Legault, Yamuna Paila, Biliana Nestorova, Baoyu Ding, Rolando Pajon, Jacqueline M Miller, Brett Leav, Andrea Carfi, Roderick McPhee, and Darin K Edwards. Preliminary analysis of safety and immunogenicity of a SARS-CoV-2 variant vaccine booster. May 2021.

30. Ching-Lin Hsieh, Jory A Goldsmith, Jeffrey M Schaub, Andrea M DiVenere, Hung-Che Kuo, Kamyab Javanmardi, Kevin C Le, Daniel Wrapp, Alison G Lee, Yutong Liu, Chia-Wei Chou, Patrick O Byrne, Christy K Hjorth, Nicole V Johnson, John Ludes-Meyers, Annalee W Nguyen, Juyeon Park, Nianshuang Wang, Dzifa Amengor, Jason J Lavinder, Gregory C Ippolito, Jennifer A Maynard, Ilya J Finkelstein, and Jason S McLellan. Structure-based design of prefusion-stabilized SARS-CoV-2 spikes. Science, 369(6510):1501–1505, September 2020.

31. Constantinos Kurt Wibmer, Frances Ayres, Tandile Hermanus, Mashudu Madzivhandila, Prudence Kgagudi, Brent Oosthuysen, Bronwen E Lambson, Tulio de Oliveira, Marion Vermeulen, Karin van der Berg, Theresa Rossouw, Michael Boswell, Veronica Ueckermann, Susan Meiring, Anne von Gottberg, Cheryl Cohen, Lynn Morris, Jinal N Bhiman, and Penny L Moore. SARS-CoV-2 501Y.V2 escapes neutralization by south african COVID-19 donor plasma. Nat. Med., March 2021.

32. Sandile Cele, Inbal Gazy, Laurelle Jackson, Shi-Hsia Hwa, Houriiyah Tegally, Gila Lustig, Jennifer Giandhari, Sureshnee Pillay, Eduan Wilkinson, Yeshnee Naidoo, Farina Karim, Yashica Ganga, Khadija Khan, Mallory Bernstein, Alejandro B Balazs, Bernadett I Gosnell, Willem Hanekom, Mahomed-Yunus S Moosa, Network for Genomic Surveillance in South Africa, COMMIT-KZN Team, Richard J Lessells, Tulio de Oliveira, and Alex Sigal. Escape of SARS-CoV-2 501Y.V2 from neutralization by convalescent plasma. Nature, March 2021.

33. Michelle M Becker, Rachel L Graham, Eric F Donaldson, Barry Rockx, Amy C Sims, Timothy Sheahan, Raymond J Pickles, Davide Corti, Robert E Johnston, Ralph S Baric, and Mark R Denison. Synthetic recombinant bat SARS-like coronavirus is infectious in cultured cells and in mice. Proc. Natl. Acad. Sci. U. S. A., 105(50):19944–19949, December 2008.

34. Victor M Corman, Olfert Landt, Marco Kaiser, Richard Molenkamp, Adam Meijer, Daniel Kw Chu, Tobias Bleicker, Sebastian Brünink, Julia Schneider, Marie Luisa Schmidt, Daphne Gjc Mulders, Bart L Haagmans, Bas van der Veer, Sharon van den Brink, Lisa Wijsman, Gabriel Goderski, Jean-Louis Romette, Joanna Ellis, Maria Zambon, Malik Peiris, Herman Goossens, Chantal Reusken, Marion Pg Koopmans, and Christian Drosten. Detection of 2019 novel coronavirus (2019-nCoV) by real-time RT-PCR. Euro Surveill., 25(3), January 2020.

35. Ali Rahimi and Benjamin Recht. Weighted sums of random kitchen sinks: replacing minimization with randomization in learning. In Nips, pages 1313–1320, 2008.

